# TYPE I INTERFERON-DEPENDENT CELLULAR REMODELING DURING ACUTE PULMONARY MHV INFECTION

**DOI:** 10.1101/2022.02.01.478627

**Authors:** S. Grabherr, A. Waltenspühl, M. Lütge, L. Büchler, H.W. Cheng, K. Zielińska, S. Caviezel-Firner, P. Krebs, B. Ludewig, N.B. Pikor

## Abstract

Impaired type I interferon (IFN) responses are predictive of severe disease during pulmonary coronavirus infection. In the periphery, dampened IFN-responsiveness is associated with viremia and hypercytokinemia, however the resolution of IFN-dependent innate immune responses in the lungs remains limited. Here, we aimed to elucidate the early dynamics of antiviral immunity and define the IFN-dependent mechanisms limiting viral spread during pulmonary infection with the murine hepatitis virus (MHV), a beta-coronavirus. While several innate immune cell types infiltrated the lungs concomitant with viral replication, the influx of type I IFN-responsive myeloid cells was essential for viral containment and prevention of fatal disease. Combining high-resolution transcriptomic analysis and genetic attenuation of interferon signaling, we delineated IFN-dependent cell-intrinsic and population-based transcriptional changes that determined viral replication and inflammatory maturation, respectively. While monocyte-derived macrophages exhibited the strongest pro-inflammatory transcriptional reprogramming during pulmonary infection, these maturation programs were impaired in the absence of IFN-signalling. Instead, IFN-deficient monocyte-derived macrophages expressed genes encoding for neutrophil attractants and IL1β, resulting in enhanced lung injury. Our results reveal the requirement for type I IFN-induced antiviral states and inflammation-induced maturation programs to secure pulmonary viral containment.

## INTRODUCTION

Coronaviruses (CoVs) are enveloped, single-stranded RNA viruses that in humans primarily cause pulmonary infections^1,2^. While the common cold CoVs are associated with mild respiratory symptoms, the emergence of pandemic strains including SARS-CoV-2 highlights the potential for CoVs to cause severe disease^3,4^. Early studies revealed elevated pro-inflammatory serum cytokine levels and dysfunctional immune responses as clinical correlates of disease severity^5-10^. Indicators of attenuated interferon (IFN) responses, including low serum levels of IFN-alpha (IFN-α) and reduced interferon-stimulated gene (ISG) signatures in peripheral blood mononuclear cells and bronchoalveolar lavage leukocytes were soon observed to be associated with severe COVID-19^11-16^. Among patients with severe disease, genetic mutations in loci responsible for type I IFN production or responsiveness^17,18^, as well as autoantibodies directed against type I IFNs^19^ may in part explain the inverse correlation between IFN-associated signatures and disease outcome. Further mechanistic insight has been offered by transcriptomic studies of peripheral blood mononuclear cells that observed the concomitant activation of NFκb signaling, which encodes a number of pro-inflammatory genes, in patients with severe COVID-19, dampened ISG signatures and viremia. Here, impaired type I IFN-mediated control of viral load, was suggested to lead to increased recognition of PAMPs and DAMPs, resulting in NFκB-mediated hypercytokinemia^11^. Nevertheless, longitudinal studies have primarily been performed on peripheral blood, and bronchoalveolar lavage sampling of incubated, intensive care patients that occurs days to weeks after initial innate IFN-mediated responses in the lungs. Thus, the cell types governing IFN-mediated viral containment in the lungs, as well as the concomitant local changes exacerbating pro-inflammatory gene signatures in the context of impaired IFN responsiveness remain elusive.

The mouse coronavirus (also known as the mouse hepatitis virus; MHV), like SARS-CoV-2, is a beta-coronavirus. Studies using MHV have elucidated key cellular players in type I IFN production and responsiveness. Plasmacytoid dendritic cells (pDCs) are the first producers of type I IFN^20^. While many cell types respond to IFN signaling, type I IFN-α receptor (IFNAR) signaling in macrophages has been shown to critically contain viral spread^21^. While IFN responsiveness in myeloid-lineage cells preserves organ function following intraperitoneal infection, the requirement of IFNAR-signaling in this cell population during pulmonary MHV infection has not been examined. Studies of mouse-adapted human coronaviruses in gene-deficient mice have revealed contradicting results for the requirements of IFNAR-signaling in controlling viral titers^22,23^, likely owing to different mutations acquired by viral passaging in different mouse strains. Alternatively, other innate immune mechanisms may account for early viral containment in the lungs. In another mucosal tissue, NK cells have been shown to limit early MHV viral spread^24^. Collectively, the cellular and molecular pathways securing immediate early viral containment during pulmonary MHV infection warrant further investigation.

Here we employed the MHV model to systematically dissect the dynamics of viral spread and anti-viral immunity in mice. We observed that the ability of myeloid cells to sense type I interferons was crucial for the survival and containment of viral load in the lungs of MHV infected mice. Single-cell RNA sequencing of lungs from infected mice revealed dominant IFN-response and pro-inflammatory cytokine gene signatures in the myeloid cell compartment. To delineate early type I IFN-dependent changes in the immunological landscape during early MHV infection, we examined the transcriptional and cellular profile of lungs from *Ifnar-*proficient and *Ifnar*-deficient LysM-Cre-targeted mice. In mice unable to sense type I interferons, we found divergent regulation of pro-inflammatory cytokines and chemokines, more influx of neutrophils, interstitial pneumonia and signs of enhanced lung injury.

## METHODS

### Mice

C57BL/6 (B6) mice were purchased from Charles River Laboratories and LysM-Cre mice were purchased from Jackson Laboratory. LysM-Cre *Ifnar*^fl/fl^ mice were provided by U. Kalinke, Helmholtz Center for Infection Research, Hannover, Germany^25^. β2M KO mice and IFNAR KO were maintained locally at the Institute for Laboratory Animal Sciences at the University of Zurich. All mice were on the C57BL/6 genetic background and were housed in the Institute of Immunobiology, Kantonsspital St. Gallen, under specific-pathogen-free conditions in individually ventilated cages. Experiments were conducted with female mice between 8 and 12 weeks of age. All experiments were performed in accordance with federal and cantonal guidelines (Tierschutzgesetz) under permission numbers SG/01/19, SG/26/20 and SG/07/21 following review and approval by the Cantonal Veterinary Office (St. Gallen, Switzerland).

### Viral infections and determination of virus titers

Mice were infected intranasally with 5×10^4^ pfu MHV-A59 or MHV-GFP as previously described^26-28^. Mice were observed daily and clinical disease symptoms and weights were noted. MHV titers were determined by standard plaque assay using L929 cells^20^.

### NK cell depletion

B6 mice were injected with 0.5 mg/ml anti-NK1.1 antibody intra-peritoneal one day prior to infection with 5×10^4^ PFU MHV. Depleting antibody was injected every second day after the first depletion.

### Isolation of hematopoietic and stromal cells

Cell isolation from mouse lungs was performed according to standardized protocols^29^. Mice were euthanized at the indicated time points and immediately perfused with PBS. Lungs were transferred into a 6-well plate containing RPMI 1640 medium (Sigma-Aldrich) supplemented with 4U/ mL elastase (Worthington), 1U/ mL Dispase II (Roche) and 10 μl/mL DNAse I (Applichem) and incubated for 45 minutes at 37°C. Lung tissue was then mechanically disrupted and washed in cold RPMI. Tissues were subject to a second enzymatic digestion in RPMI 1640 medium supplemented with 25 μg/mL Liberase TM (Roche) and 0.25 μg/mL DNAse I and incubated for 30 minutes at 37 °C. Any larger tissue fragments were dissociated mechanically followed by washing of the cell strainer with MACS buffer (PBS, 2% FCS, 2mM EDTA (Sigma-Aldrich)). Stromal and hematopoietic cell fractions were enriched using MACS anti-CD45 microbeads (Miltenyi Biotec).

### Flow cytometry

Cell characterization was performed by incubating single-cell suspensions in PBS containing 2% FCS and 2 mM EDTA for 20 min at 4 °C, using the indicated antibodies (Supplementary Table 1). Ghost Dye Violet 510 (LubioScience) was used to identify dead cells. Antigen-specific T cells were detected using PE-conjugated s598 tetramers (Sanquin Reagents) after an incubation of 20 min at 37 °C. For measuring cytokine production, 10^6^ cells were incubated with brefeldin A (5 μg/ml; Sigma) for 3.5 h at 37 °C. For intracellular staining, cells were surface stained and fixed using Cytofix/Cytoperm (BD Biosciences) for 20 min. Fixed cells were incubated at 4 °C for 40 min with monoclonal anti-GzmB (eBioscience) diluted in permeabilization buffer (eBioscience). Samples were analyzed by flow cytometry using a FACS Fortessa flow cytometer and the FACSDiva operating program. Data were analyzed using FlowJo software (Tree Star).

### Single-cell RNA sequencing analysis

Cell sorting was performed on a BD FACSMelody™ cell sorter using BDChorus software. For discrimination between live and dead cells, Seven-aminoactinomycin D (Sigma-Aldrich) was added prior to sorting. Sorted cells were passed through the 10x Chromium (10x Genomics) system and cDNA libraries were generated according to the manufacturer’s recommendations (Chromium Single Cell 3′ Reagent Kit (version 3 chemistry)). Sequencing of libraries was done using the NovaSeq 6000 Illumina sequencing system at the Functional Genomic Center Zürich. Initial processing and gene expression estimation were performed using Cell Ranger (version 5.0.1) with the Ensembl GRCm38.94 release as a reference, as well as the MHV-A59 Nucleocapsid gene sequence (NC_001846.1) to build index files for alignments. Subsequent quality control was performed with the scater (version 1.20.1)^30^ and SingleCellExperiment (v1.10.1)^31^ R/Bioconductor packages in R version 4.0.1. Cells with particularly high or low numbers of detected genes or UMI counts (more than two median absolute deviations from the median across all cells) as well as cells with a large fraction of mitochondrial genes (more than two median absolute deviations above the median across all cells) were removed to avoid contamination with damaged cells. After quality control and removal of contaminants, in total 8196 cells from naïve B6 samples, 14646 cells from infected B6 controls (7558 macrophages) and 12774 cells from infected LysM-Cre *Ifnar*^fl/fl^ samples (2434 macrophages) were retained for further processing using the Seurat package (version 4.0.1)^32^. Data were analyzed using functions from the Seurat package for normalization, scaling, dimensional reduction with principal component analysis (PCA) and tSNE, and graph-based clustering. Resulting clusters were characterized based on expression pattern of canonical marker genes and cluster markers calculated with the FindMarkers function of the Seurat package. To compare expression profiles across conditions differentially expressed genes were conceived from Wilcoxon test as implemented in the FindMarkers function. To investigate functional differences between inflammatory macrophages in LysM-Cre *Ifnar*^fl/fl^ and control mice, differentially expressed genes in macrophage clusters were tested for enrichment in gene ontologies using the ‘enrichGO’ function from the clusterProfiler R/Bioconductor package (version 3.18.1)^33^. Top significant ontologies (q value < 0.05) were visualized. Functional gene signatures were summarized from top differentially expressed genes based on their reported functions and projected on a diffusion map^34^ by running the ‘DiffusionMap’ function from the destiny R package (3.4.0)^35^.

### Immunohistochemistry

Murine lungs were perfused and fixed overnight at 4 °C in 4% paraformaldehyde (Merck Millipore) under agitation. Tissues were embedded and oriented in 4% low-melting agarose (Invitrogen) in PBS and cut into 60 to 80 um sections using a vibratome (VT-1200, Leica). Sections were blocked in PBS containing 10% FCS, 1 mg/ml anti-Fcγ receptor (BD Biosciences) and 0.1% Triton X-100 (Sigma). Sections were stained overnight with the indicated antibodies as indicated in Supplementary Table 1. Unconjugated antibodies were detected with the following secondary antibody incubation for 1 hr at room temperature. Nuclei were stained with 4′-6-diamidino-2-phenylindole (DAPI) dihydrochloride (Life Technologies), and lung sections were mounted on glass microscopy slides using fluorescence mounting medium (Dako). Imaging was performed with an LSM 980 confocal microscope (Carl Zeiss) and microscopy data were processed with ZEN software (Zeiss) and Imaris version 9 software (Bitplane).

### *In vitro* infections

Bone marrow was isolated from bones from IFNAR KO mice, LysM-Cre *Ifnar*^fl/fl^ mice and littermate controls. Murine bone marrow–derived macrophages (BMDMs) were generated by 7 days of culture with granulocyte-monocyte colony-stimulating factor containing supernatant. After 7 days, 10^5^ BMDMs were plated into 48 well plates and on the following day incubated with 0.01 MOI of MHV at 37 C. After 1 hour the virus was replaced with medium and supernatant was collected after 6 or 12 hours. MHV titers in the supernatant were determined by standard plaque assay.

### Cytokine validations (CBA and ELISA)

For cytokine measurements, lung supernatant of day 2 infected mice and uninfected mice were used. IL1β levels were measured using the IL1β ELISA kit (Invitrogen) according to manufacturer instruction. Absorbance was measured at 450nm on a Microplate reader (Tecan).

TNF-α, MCP-1 and IL-6 cytokine levels were quantified using the Cytometric bead assay Mouse inflammation kit (BD Bioscience) as recommended by the manufacturer and measured by flow cytometry using a FACS Fortessa flow cytometer.

### Vascular permeability

Vascular permeability assays were performed as per standardized protocol^36^. 40kDa FITC-Dextran (Invitrogen) was injected intravenously into mice. 1 hour later, mice were sacrificed and blood was collected by cardiac puncture. Serum was diluted 1:10 in PBS, and the fluorescence intensity was measured at 528 nm on a Microplate reader (Tecan).

### Statistical analyses

Statistical analyses were performed with GraphPad Prism 8.0 with Mann-Whitney test and longitudinal comparison between different groups was performed with one or two-way analysis of variance (ANOVA) with Tukey’s post-test or Kruskal Wallis test with Dunn’s post-test. Statistical analyses are indicated in the figure legends. Statistical significance was defined as P < 0.05.

## RESULTS

### Type I IFNs critically control acute pulmonary MHV infection

To systematically assess viral spread along the respiratory tract, we infected C57BL/6 (B6) mice with MHV via the intranasal (i.n.) route and measured viral titers between day 0 and day 6. All mice used for viral titer analysis showed presence of virus in at least one organ (Supplementary Figure 1A). Intranasal infection of mice led to detectable viral titers in the NALT (Nasopharyngal-associated lymphoid tissue), the trachea and lung already 12 hours post infection (p.i.) (Figure 1A). Viral titers in the lungs peaked between 24 and 48 hours after the infection. By day 6 p.i. most mice had cleared the virus in the upper and lower respiratory tract. In cervical lymph nodes (cLN) viral titers were detectable from 24 hours after the infection with complete clearance by day 6. Clusters of virus-infected cells were clearly visible in the lungs of MHV-infected mice at day 2 p.i. (Figure 1B). Viral infection was accompanied by an influx of myeloid cells, especially inflammatory monocytes/macrophages (Figure 1C, Supplementary Figure 1 B,C) and Granzyme B (GzmB) expressing NK cells by day 2 p.i. (Figure 1D; Supplementary Figure 1D,E). By day 8 post infection, the number of virus-specific s598-tetramer binding CD8^+^ and activated KLRG1^+^ CD62L^-^ CD8^+^ T cells was observed to peak in the lung (Figure 1E,F; Supplementary Figure 1F,G). Previous work has demonstrated the requirement for type I IFN responsiveness in myeloid cells, as well as a role for NK cells in controlling acute MHV infection^24^. Given the rapid increase in inflammatory macrophages and GzmB^+^ NK cells observed in the lungs of MHV-infected mice, we sought to delineate the relevant cell types mediating viral containment during pulmonary MHV infection. To this end, we assessed the survival of mice lacking the gene encoding the interferon alpha receptor (*Ifnar*) in LysM-Cre targeted myeloid cells, as well as in mice treated with the NK-depleting α-NK1.1 antibody. Consistent with peripheral MHV infection^26^, mice lacking IFNAR in myeloid-lineage cells had worse survival compared to infected Cre^-^ littermate controls (Figure 1G), while NK cell depletion was not observed to impact mouse survival (Figure 1H). As demonstrated previously^27^, the presence of CD8^+^ T cells was crucial for survival of mice, as CD8^+^ deficient β2-Microglobulin (β2M) mice were observed to succumb to the disease between day 8 and 10 p.i. (Figure 1I). Taken together, pulmonary MHV infection was accompanied by increased inflammatory macrophage and NK cell infiltration, although type I interferon responsiveness in myeloid cells crucially was crucial to control MHV infection.

**FIGURE 1.**
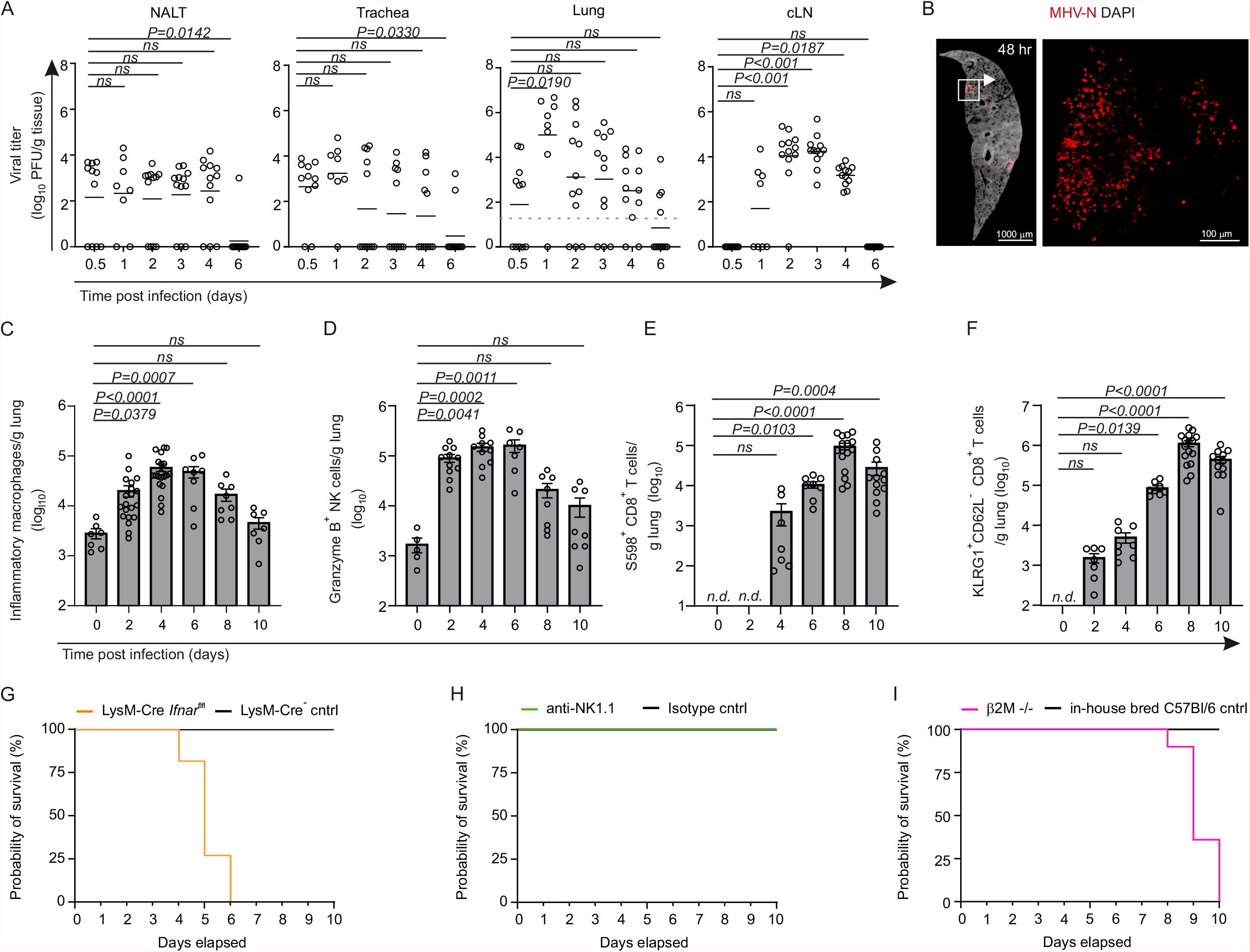
A) Viral titer of nasal associated lymphoid tissue (NALT), trachea, lung and cervical LN (cLN) over time in mice infected intranasally with 5×10^4^ PFU MHV. B) Representative immunofluorescence images of lungs showing MHV-nucleocapsid staining in infected cell clusters in intranasally-infected B6 mice 48hrs post infection (p.i.). Scale bar 1000 μm (overview) and 100 μm (boxed area). C-F) Flow cytometric cell enumeration of C) inflammatory macrophages, D) Granzyme B^+^ NK cells, E) s598^+^ CD8^+^ T cells and F) KLRG1^+^ CD62L^-^ CD8^+^ T cells of MHV-infected B6 mice at the indicated days after infection. G-I) Survival curves following MHV infection of: G) LysM-Cre *Ifnar*^fl/fl^ mice compared to LysM-Cre-negative controls, H) anti-NK1.1 depleted B6 mice compared to IgG isotype-treated controls, and I) b2M^-/-^ compared to in-house bred B6 mice. Values indicate mean ± s.e.m. for each time point or organ analysed. (A) Pooled data from three independent experiments with NALT and Trachea N=11 (day 0.5), N=12 (days 1-6); cLN N=10 (day 0.5), N=12 (days 1-6); Lung N=11 (day 0.5), N=8 (day 1), N=12 (days 1-6). (B) Image is representative of at least 6 mice on day 2 following MHV infection. (C) Pooled data from two-three independent experiments with N=7 (d0, d7), N=20 (d2, d4), N=8 (d6, d8). (D-F) Pooled data from two-three independent experiments with N=10 (d0), N=8 (d2, d6), N=7 (d4), N=16 (d8) and N=12 (d10). In (E-F) T cell counts are normalized to day 0. (G) Two experiments with N=9 in each group (H) One experiment with N=4 in each group (I) One experiment with N=5 in each group. Statistical analysis was performed using Kruskal-Wallis test with Dunn’s multiple comparisons test.

### MHV infects and transcriptionally reprograms multiple cell types in the lung

At present, insight into the inflammation-induced remodeling of coronavirus-infected lungs is primarily limited to sampling of bronchoalveolar lavage at later stages of viral infection, or to post-mortem tissue analysis. To define the early cellular and molecular changes in the lungs during MHV infection, we sought to examine the transcriptomic landscape of lung tissue from infected compared to naïve mice. We anticipated that virus-infected cells would account for a small percentage of overall lung cells. To enrich for infected cells we infection mice with the fluorescent MHV-GFP virus. Indeed, the proportion of infected cells 36 hours after the infection was around 0.3% of live cells (Figure 2A,B). To characterize the hematopoietic and non-hematopoietic compartments of lung tissue, we sorted for CD45^+^/CD45^-^ cells from naïve mice, and for GFP^+^ CD45^+^/CD45^-^ and GFP^-^ CD45^+^/CD45^-^ cells from MHV-GFP infected lungs (Figure 2C). Cell types were assigned based on *Sftpc, Krt8, Epcam* and *Muc 1* for epithelial cells; *Tagln, Col3a1, Pdgfra* and *Pdgfrb* for fibroblasts; *Cdh5, Pecam1, Cldn5, Car4, Ednrb* and *Apln* for Aerocytes; *Plvap, Cd93* and *Gpihbp1* for Capillaries; *Fabp1, Ear1, Mrc1, Marco, Gpnmb, Ear1* and *Siglecf* for alveolar macrophages; *Fcgr4, Treml4, Fabp4* and *Cx3cr1* for interstitial macrophages; *Itgam, Aif1, Ccr2* and *Ly6c* for monocyte-derived macrophages; *Batf3, Siglech* and *H2-Ab1* for DC-pDCs; *S100a9, Ly6g, Retnlg* and *Slpi* for neutrophils; *Ncr1, Nkg7* and *GzmB* for NK cells; *Cd3e, Cd3g, Il7r* and *Trbc2* for T cells; and *Cd79a, Cd19* and *Igkc* for B cells (Supplementary Figure 2A)^37-40^. To identify virus-infected cells, we examined the expression of MHV Nucleocapsid transcripts, which were broadly detected in myeloid cell populations, epithelial cells and lung fibroblasts and endothelial cells (Figure 2E,F). The presence of viral Nucleocapsid protein was confirmed by histological staining of lung tissue. Nucleocapsid staining was observed in CD45^+^ hematopoietic cells (Figure 2G), Surfactant protein C (SFTPC)-expressing alveolar type II epithelial cells (Figure 2H), and Podoplanin (PDPN)^+^ stromal cells (Figure 2I). In addition, we could further validate the presence broadly-infected cell types in the hematopoietic and stromal cell compartment in the lungs by flow cytometric analysis (Supplementary Figure 2C,D). Monocyte-derived macrophages were the most infected cell type in the hematopoietic compartment (Supplementary Figure 2C), whereas blood endothelial cells, fibroblasts and alveolar-epithelial cells type II seem to be to most viral-targeted cell types in the stromal cell populations (Supplementary Figure 2D).

**FIGURE 2.**
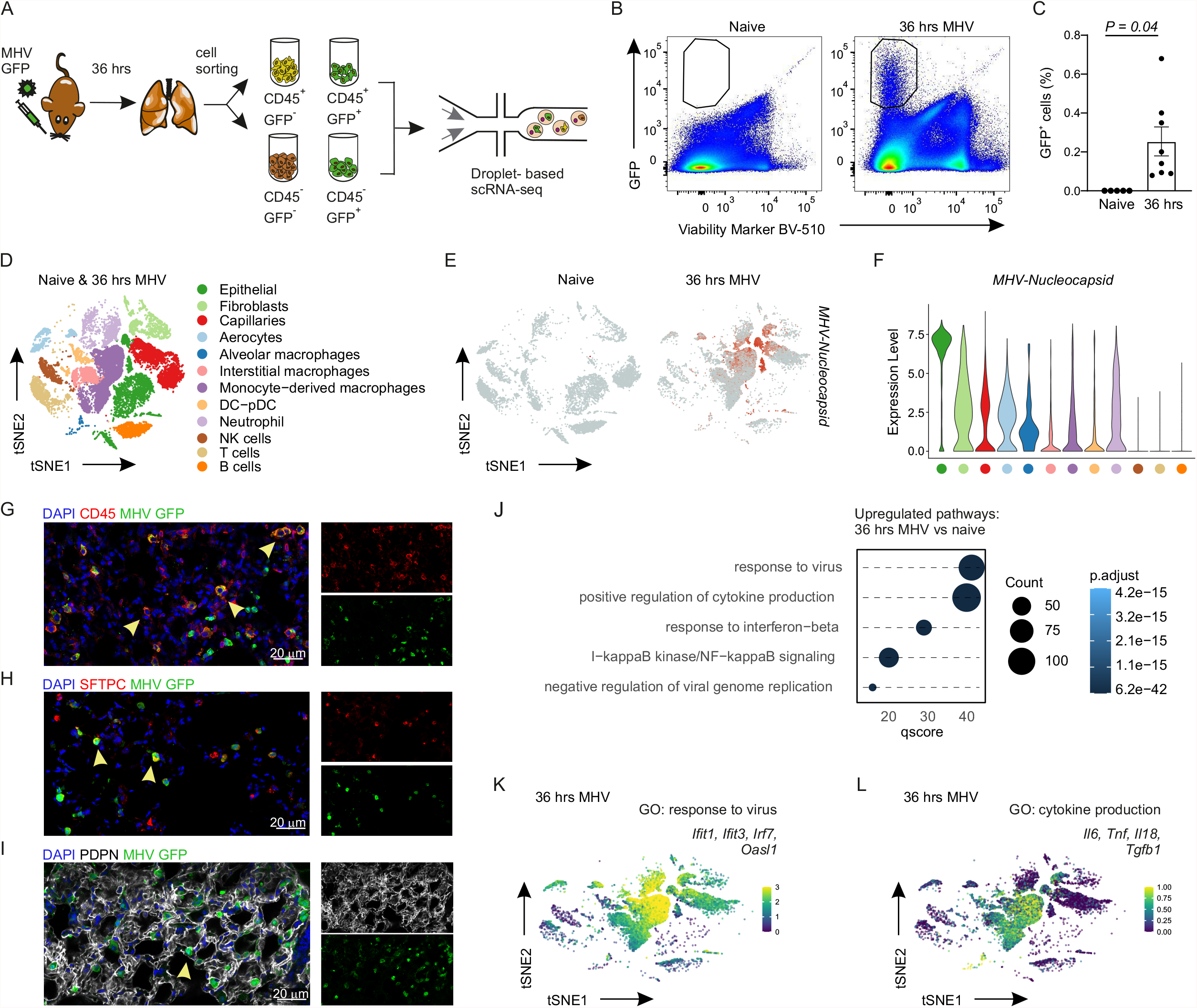
A) Schematic representation of the experimental approach from cell isolation to droplet-based single-cell RNA sequencing. B) Representative flow cytometric plots showing the presence of GFP^+^ cells in the lungs of infected but not naive mice at 36h p.i with MHV-GFP. C) Flow cytometric analysis of GFP^+^ cells in naïve and 36h p.i. mouse lungs D) Dimensional reduction t-SNE plots demonstrating the cell types in the naive and 36 hr-infected lungs based on marker gene expression. E) Feature plot representing the expression pattern of the MHV Nucleocapsid transcript. F) Violin plot of Nucleocapsid transcript levels across cell types. G-I) Representative high-resolution immunofluorescence images of G) hematopoietic CD45^+^ cells H) SFTPC^+^alveolar epithelial type II cells I) PDPN^+^ stromal cells. Scale bars 20 μm. J) Significantly enriched gene ontologies among all cell types in the lungs of B6 mice 36 hrs p.i. compared to naïve mice. K-L) t-SNE plots of 36hr-infected B6 mice with projected genes of the indicated GO pathways. (C) Statistical analysis was performed using unpaired two-tailed Student’s t-test.

Given the breadth of infected cell types, we sought to assess the transcriptional programs that were differentially regulated across cell types of MHV-infected lungs. Gene set enrichment analysis (GSEA) of signaling pathways enriched in infected compared to naïve mice revealed an upregulation of pathways associated with anti-viral defense mechanisms, such as regulation of innate immune response, pattern recognition receptor signaling or cytokine signaling (Figure 2J). While interferon stimulated gene (ISG)-signatures were enhanced across infected cell types in MHV-infected lungs, macrophages and neutrophils cells harbored a strong ISG-signature (Figure 2K), while transcripts of pro-inflammatory cytokines were preferentially expressed by monocyte-derived macrophages (Figure 2L). In sum, although MHV was observed to readily infect hematopoietic, epithelial and stromal cell types, IFN-responsive and pro-inflammatory transcriptional programs were most pronounced in the monocyte-derived macrophages.

### Impaired IFNAR responsiveness in myeloid cells permits increased viral replication and host cell death

Since sensing of type I interferons by myeloid cells was observed to secure mouse survival following pulmonary MHV infection, we sought to determine the cellular and molecular transcriptional changes that accompany impaired IFN responsiveness. To this end, we performed scRNA-seq of lungs from infected LysM-Cre *Ifnar*^*fl/fl*^ mice 36 hours p.i. to reveal changes on the transcriptional level (Figure 3A). MHV-N transcripts were expressed in the same cell populations as in wildtype mice, indicating a similar viral tropism in the conditional absence of IFNAR. Nevertheless, the proportion of infected cells appeared to be reduced compared to IFNAR-sufficient controls. (Figure 3B). Relative flow cytometric quantification of infected cells using MHV-GFP confirmed a reduction in the number of infected cells in mice lacking IFNAR in LysM-Cre-targeted myeloid cells (Figure 3C,D). As expected, cells from the lungs of LysM-Cre *Ifnar*^*fl/fl*^ conditional knock out mice revealed an impaired type I interferon signalling signature compared to controls (Figure 3E). In addition, GSEA analysis of lung macrophages revealed the failure of cells in the lungs of infected LysM-Cre *Ifnar*^*fl/fl*^ mice to downregulate ribosomal and translational pathways compared to controls (Figure 3F), suggesting that infected cells lacking IFNAR may be better able to support viral replication and new virion assembly. Indeed, bone-marrow derived macrophages from LysM-Cre *Ifnar*^*fl/fl*^ mice showed more efficient viral replication and reduced viability 12 hours after *in vitro* infection with MHV compared to controls (Figure 3G,H). Consistent with elevated viral titers observed *in vitro*, mice unable to respond to type I IFN by myeloid cells showed significantly impaired viral clearance in the lungs at day 4 p.i. (Figure 3I). The presence of elevated viral titers in the lungs of LysM-Cre *Ifnar*^fl/fl^ mice was accompanied by a concomitant increase in CD11b^+^ CD11c^+^ cell numbers in the lung (Figure 3J). In sum, impairing the sensing of type I interferon by myeloid cells was observed to lead to impaired induction of ISG-regulated antiviral programs at a single cell-level, resulting in decreased survival of infected host cells and impaired early viral control in lungs of infected mice.

**FIGURE 3.**
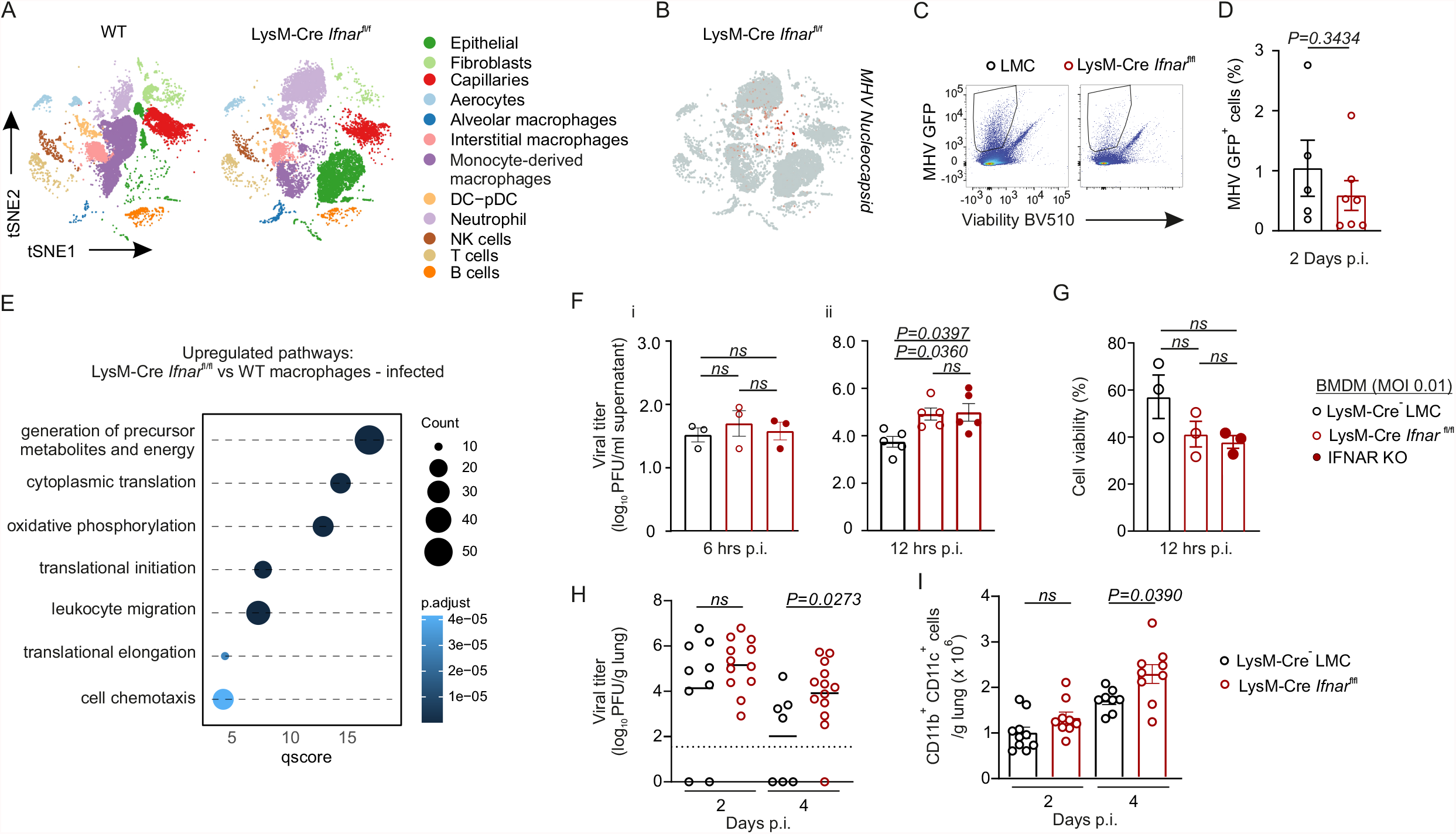
A) Dimensional reduction t-SNE plot of pulmonary cell populations from LysM-Cre *Ifnar*^fl/fl^ and WT mice 36h post MHV infection. B) Feature plot representing the expression pattern of the MHV Nucleocapsid transcript in the lungs of infected LysM-Cre *Ifnar*^fl/fl^ mice. C) Representative flow cytometric plots of GFP^+^ cells from with MHV-GFP. D) Quantification of the percent abundance of GFP^+^ cells in the lungs of LysM-Cre *Ifnar*^fl/fl^ and littermate control (LMC) infected mice. E) Significantly enriched gene ontologies at 36 hours post-infection in lung macrophages of LysM-Cre *Ifnar*^fl/fl^ compared to WT mice. F) Viral titers from viral supernatant of bone-marrow derived macrophages of *Ifnar*-/, LysM-Cre *Ifnar*^fl/fl^ and LMC mice *in vitro* at the indicated time points after infection (MOI 0.1) with MHV. G) Cell viability of bone-marrow derived macrophages 12h after infection with MHV *in vitro*. H) Viral titers in the lungs of LysM-Cre *Ifnar*^fl/fl^ and LMC mice at days 2 and 4 after intranasal infection with 5×10^4^ PFU MHV. I) Absolute cell numbers of CD11b^+^ CD11c^+^ myeloid cells in the lungs of LysM-Cre *Ifnar*^fl/fl^ and LMC mice at days 2 and 4 after MHV infection. Values indicate mean ± s.e.m. for each time point or organ analysed. (D) Pooled data from two independent experiments with N=5 (LMC) and N= 7 (CKO). (F) Pooled data from (i) two independent experiments with N= 3 and (ii) from three independent experiments with N=5. (G) Pooled data from two independent experiments with N= 3. H) Pooled data from three independent experiments with N= 9 (LMC d2), N=12 (CKO d2, CKO d4), N=7 (LMC d4). (I) Pooled data from three independent experiments with N= 10 (LMC d2), N=9 (CKO d2, CKO d4), N=8 (LMC d4). (F), (G), (I) P values as per one-way ANOVA with Tukey’s multiple comparisons test. (H) Statistical analysis was performed using unpaired two-tailed Student’s t-test.

### IFN-dependent, infection-associated maturation of monocyte-derived macrophages

To further determine the subset identity of macrophages from *Ifnar*-proficient and *Ifnar*-deficient LysM-Cre-targeted cells we re-embedded monocyte-derived, interstitial and alveolar macrophage cell types from naïve, *Ifnar*-proficient and *Ifnar*-deficient mice. While macrophage subset identity was maintained in each condition, dimensional reduction plots revealed little overlap between cells from different gentoypes or from uninfected lungs (Figure 4A), suggesting a change in activation state. We used diffusion maps to plot macrophages along their activation trajectories, and found an interferon-dependent, infection-induced transcriptional reprograming of monocyte-derived macrophages (Figure 4B). While a modest transcriptional remodeling was observed in interstitial macrophages, too few alveolar macrophages were captured to assess the extent of inflammation-associated changes. Gene set enrichment analysis of *Ifnar*-proficient macrophages from MHV-infected mice revealed the upregulated expression of inflammatory and antiviral programs, including response to virus, response to interferon-beta, cytokine-mediated and NFκB signaling compared to *Ifnar*-deficient macrophages (Figure 4C). Analysis of cytokine and chemokine expression profiles revealed an upregulation of IFN-stimulated chemokines such as *Cxcl10*, but also pro-inflammatory cytokines including *Il6* and *Tnf*, especially by monocyte-derived macrophages (Figure 4D). Consistent with the observed enrichment of genes associated with cell chemotaxis in *Ifnar*-deficient macrophages (Figure 3E), we also observed the enhanced transcript expression of *Il1b*, the alarmins *S100a8* and *S100a9*, as well as *Ccl6* and *Mif* (Figure 4D). The contribution of macrophage-associated gene expression changes to the protein cytokine levels in the lungs was assessed using ELISAs and cytokine bead assays of lung supernatant from mice infected with MHV two days earlier. Consistent with transcriptome-level gene expression, the lungs of LysM-Cre *Ifnar*^fl/fl^ mice showed a significant increase in activated IL-1β compared to naïve mice (Figure 4E). In contrast, lungs of conditionally *Ifnar*-deficient mice demonstrated significantly less MCP1 (encoded by *Ccl2*) and a trend towards less TNFα?compared to *Ifnar*-proficient mice (Figure 4F, G). Collectively, these data reveal a dynamic IFN-dependent, infection-associated transcriptional reprogramming of monocyte-derived macrophages, that in the absence of *Ifnar* compensates with increased IL1β expression and an altered chemotactic and alarmin profile.

**FIGURE 4.**
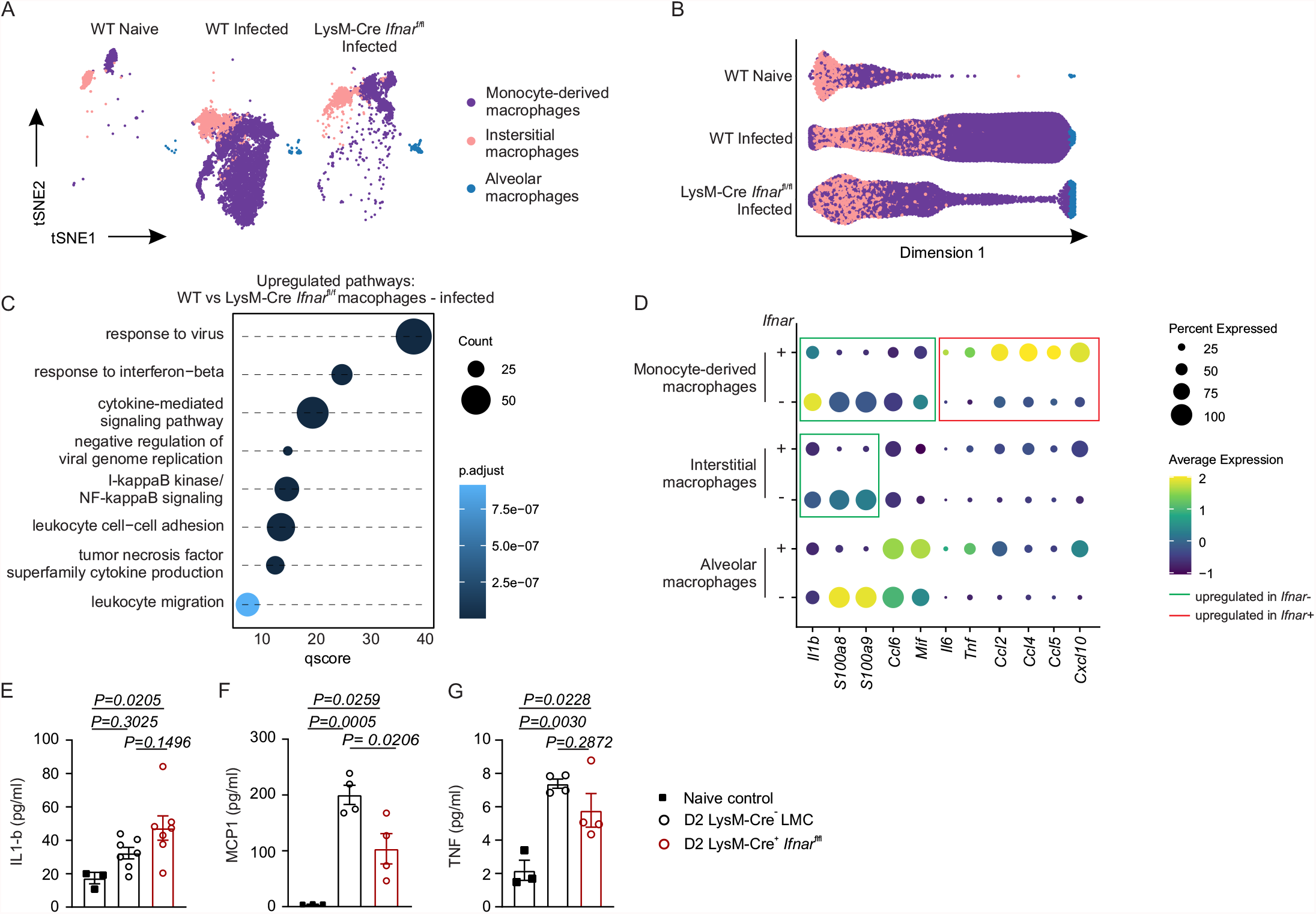
A) Dimensional reduction t-SNE plot of the indicated macrophages populations in the lung of Naïve WT, Infected WT and of LysM-Cre *Ifnar*^fl/fl^ mice. B) Diffusion map showing subset clustering of naïve WT, infected WT and of infected LysM-Cre *Ifnar*^fl/fl^ macrophage populations. C) Significantly enriched gene ontologies at 36 hours post-infection in lung macrophages of WT mice compared to LysM-Cre *Ifnar*^fl/fl^ mice. D) Average expression of inflammatory cytokines in macrophage populations sufficient and deficient of *Ifnar* in myeloid cells. E-G) Cytokine levels in the lung supernatant of LysM-Cre *Ifnar*^fl/fl^ mice compared to LMC and naïve mice. E) IL1-β levels obtained by ELISA; data is representative of two independent experiments. F-G) CBA readout of F) MCP1 and G) TNF in the lungs of infected mice; data is representative of two independent experiments. (E) – (G) P values as per one-way ANOVA with Tukey’s multiple comparisons test.

### Conditional *Ifnar* deficiency exacerbates pulmonary inflammation and vascular permeability

Given the transcriptional and protein-level changes in inflammatory mediators in the lungs of *Ifnar*-proficient and *Ifnar*-deficient LysM-Cre-targeted mice, we next sought to investigate whether the altered inflammatory profile impacted the pulmonary immune cell composition of MHV-infected mice. Flow cytometric analysis of myeloid cells in the lungs revealed that the numbers of alveolar and interstitial macrophages were significantly increased in LysM-Cre *Ifnar*^*fl/fl*^ mice on day 4 p.i. (Figure 5A,B, Supplementary Figure 3A), while inflammatory monocytes/macrophages only show a trend of an increase in conditionally gene-deficient mice (Figure 5C, Supplementary Figure 3B). In contrast, the absolute number of neutrophils was observed to be significantly increased in the lungs of LysM-Cre *Ifnar*^*fl/fl*^ mice compared to littermate controls already at day 2 p.i. and remained increased at day 4 (Figure 5D, Supplementary Figure 3C). Consistent with increased pulmonary inflammation, LysM-Cre *Ifnar*^*fl/fl*^ showed significantly higher vascular permeability as measured by serum FITC-Dextran fluorescence intensity compared to uninfected controls on day 2 following compared to littermate controls (Figure 5E). Taken together, a selective impairment of type I interferon responsiveness in lungs of MHV-infected mice was observed to culminate in increased early neutrophil inflammation, increased vascular leakage and subsequent interstitial inflammation.

**FIGURE 5.**
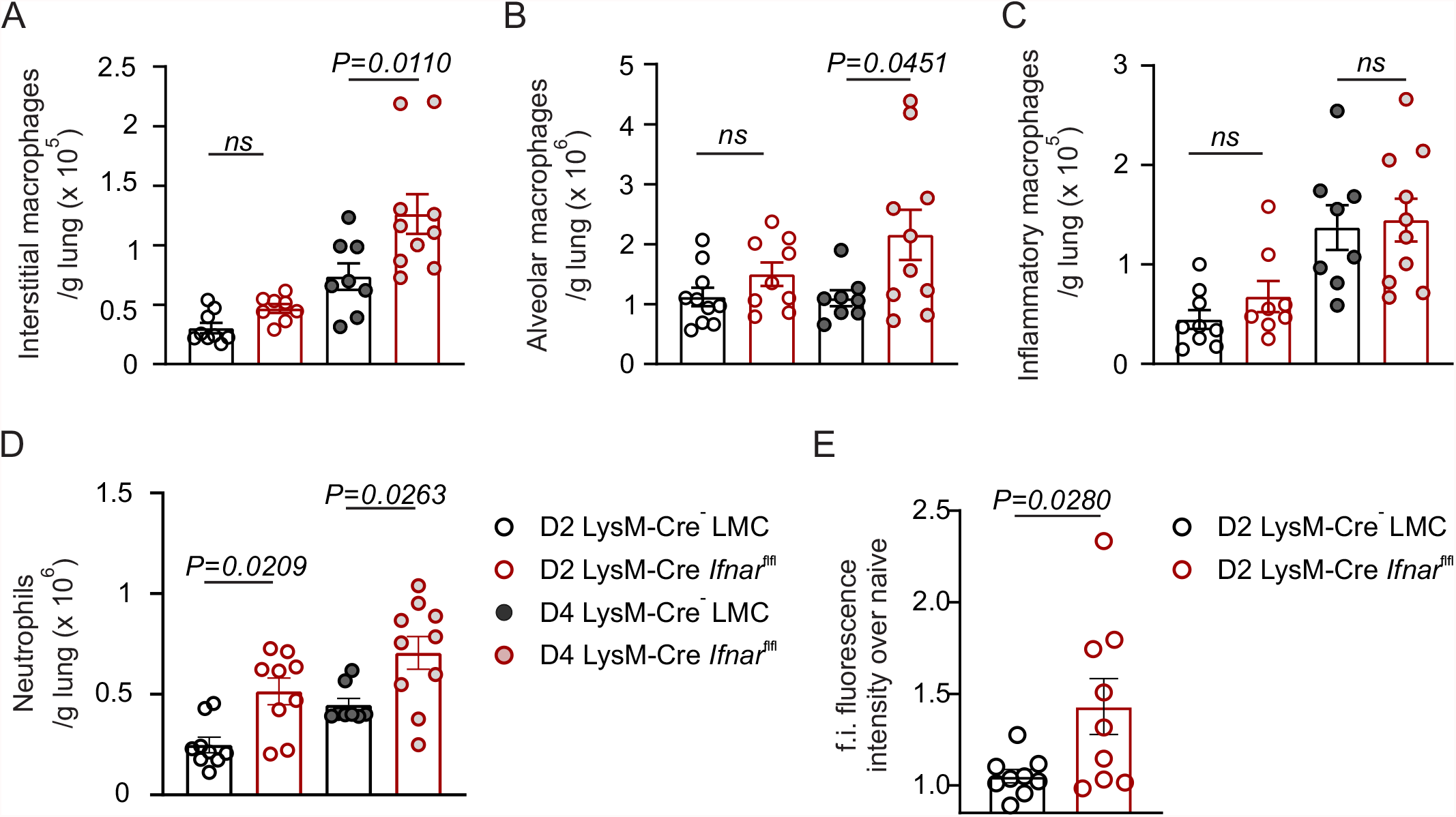
A-D) Flow cytometric cell enumeration of A) interstitial macrophages, B) alveolar macrophages, C) inflammatory macrophages and D) neutrophils of MHV-infected LysM-Cre *Ifnar*^fl/fl^ mice compared to LMC at the indicated time points after infection. E) Fluorescence intensity from serum of LysM-Cre *Ifnar*^fl/fl^ and LMC mice previously injected with FITC-Dextran intravenously as an indication of vascular permeability. (A-D) Pooled data from three independent experiments with (A) N= 9 (LMC d2), N=8 (CKO d2, LMC d4), N=10 (CKO d4); (B) N= 10 (LMC d2), N=9 (CKO d2, LMC d4), N=10 (CKO d4); (C)) N= 9 (LMC d2), N=8 (CKO d2, LMC d4), N=10 (CKO d4); (D) N= 9 (LMC d2, CKO d2), N=8 (LMC d4), N=10 (CKO d4). (E) Pooled data from three independent experiments with N= 9 (LMC d2) and N=10 (CKO d2). (A) -(D) P values as per one-way ANOVA with Tukey’s multiple comparisons test. (E) Statistical analysis was performed using unpaired two-tailed Student’s t-test.

## DISCUSSION

Type I interferons are early-induced cytokines that limit various stages of the viral lifecycle and promote an antiviral immunity at the site of infection^41^. Coronaviruses are highly pathogenic viruses that require type I interferon responses to control viral infection, with attenuated interferon IFN responsiveness being associated with exacerbated disease severity following SARS-CoV-2 infection^11-16^. While many high-dimensional analyses have profiled the dysregulated immune responses in peripheral blood or bronchoalveolar lavage that correlate with dampened type I IFN signals, many of these analyses occur long after the induction of interferons immediately following viral infection, leaving the local, early type I IFN-dependent antiviral and inflammatory changes in the lungs following coronavirus infection unresolved. Here, we employed a pulmonary infection model instigated by the murine hepatitis virus (MHV), a mouse beta-coronavirus, to dissect the early inflammation-induced and type I interferon-dependent transcriptional programs in the lungs of infected mice. Our findings reveal a profound early accumulation of innate immune cells types into the lungs within hours of detectable viral replication in the lungs. While inflammatory macrophages, neutrophils and activated NK cells were found to accumulate in infected lungs, uniquely IFN-responsive macrophages critically controlled viral infection, preventing fatal disease. Monocyte-derived macrophages were the main producers of pro-inflammatory cytokines in the lungs following infection, but also as the primary macrophage cell type preferentially infected by MHV. In the absence of IFNAR-induced signaling cascades, lung-infiltrating, monocyte-derived macrophages were highly susceptible to viral infection, dampening overall levels of proinflammatory mediators and allowing uncontrolled viral replication in the lung. These events were compensated by increased IL-1β and neutrophil chemoattraction, accompanied by increased vascular permeability and lung injury. Collectively, our results delineate the IFN-dependent responses in the myeloid cell types in the lung during early coronavirus infection.

Several studies of murine coronavirus infection point to an unclear role for type I IFNs in limiting viral replication. Studies of mouse-adapted human coronaviruses in gene-deficient mice have revealed contradicting results for the requirements of IFNAR-signaling in controlling viral titers^22,23^. In mice infected with mouse-adapted SARS-CoV, delayed type I IFN responses exacerbated severe coronavirus infection revealing a temporal requirement for protective type I IFN-mediated innate immunity^22^, while another study of IFNAR-deficient mice revealed little change in viral titers^23^. Our temporal analysis of early MHV infection using high dimensional transcriptomic analysis, genetic ablation of type I IFN signaling and *in vitro* experiments, reveals a highly dynamic role for type I IFN-dependent antiviral and inflammatory changes in the lungs during cytopathic viral infection. On a per-cell level, IFNAR signaling induces the anticipated antiviral programs that limit viral replication; MHV replicates more quickly in, yet reduces the survival of infected *Ifnar*-deficient cells. While we observed only a trend towards reduced infected cell numbers in the lungs using a fluorescent MHV virus, in vivo quantification of replicating virus and leukocyte numbers in the lungs, revealed the need for sufficient myeloid cell influx to culminate in an appreciable increase in viral titers. The preferential infectively of pulmonary macrophages is consistent with previous studies identifying macrophages as an important sink for MHV infection^21^. In the case of pulmonary MHV infection, when this sink is depleted in the absence of IFNAR-induced antiviral programs, the primary cell population producing pro-inflammatory mediators that likely act as a second-line innate immune defense to contain virus is also gone, licensing further uncontrolled viral replication. These events culminate in increased neutrophil accumulation in the lungs, which was associated with increased vascular permeability, creating a means for systemic viral spread. It is then the multi-organ infection with MHV that culminates in organ failure and fatal disease^26^.

Distinct coronavirus strains evolve different immune evasion strategies and cellular tropisms that co-evolve to take advantage of a plethora of host proteins beyond the host cell entry receptor^42^. Here, we used a native mouse coronavirus to dissect the cellular and molecular changes that occur in the lungs over the course of acute coronavirus infection. Our results reveal that early type I interferon-dependent survival of inflammation-matured cell types is critical to mount a sufficient, early antiviral immune response and efficiently contain viral infections. In the context of early pulmonary coronavirus infection, impaired IFN responsiveness culminates in profuse viral replication, increased neutrophilia and vascular injury and culminates in systemic spread and fatal infection.

## SUPPLEMENTARY TABLES

**TABLE 1:** Antibodies for flow cytometric analysis.

| **Antibody** | **Clone** | **Fluorochrome** | **Cat. number** | **Supplier** |
| --- | --- | --- | --- | --- |
| anti-Ly6C | HK1.4 | PerCP | 128028 | BioLegend |
| anti-CD11c | N418 | APC-Cy7 | 117324 | BioLegend |
| anti-I-A/I-E | M5/114.15.2 | AF700 | 107622 | BioLegend |
| anti-CD24 | M1/69 | Al. Fluor 647 | 101818 | BioLegend |
| anti-CD11b | M1/70 | BV711 | 101242 | BioLegend |
| anti-CD45 | 104 | BV605 | 109841 | BioLegend |
| anti-CCR2 | SA203G11 | BV421 | 150605 | BioLegend |
| anti-CD64 | X54-5/7.1 | PE-Cy7 | 139314 | BioLegend |
| anti-Ly6G | 1A8 | PE | 551461 | BD Biosciences |
| anti-CD31 | 390 | PerCP | 102420 | BioLegend |
| anti-CD45 | 30-F11 | APC-Cy7 | 103116 | BioLegend |
| anti-CD24 | M1/69 | Al. Fluor 647 | 101818 | BioLegend |
| anti-I-A/I-E | M5/114.15.2 | BV421 | 107632 | BioLegend |
| anti-CD326 | G8.8 | PE-Cy7 | 118216 | BioLegend |
| anti-PDPN | 8.1.1 | PE | 127408 | BioLegend |

**TABLE 2:**
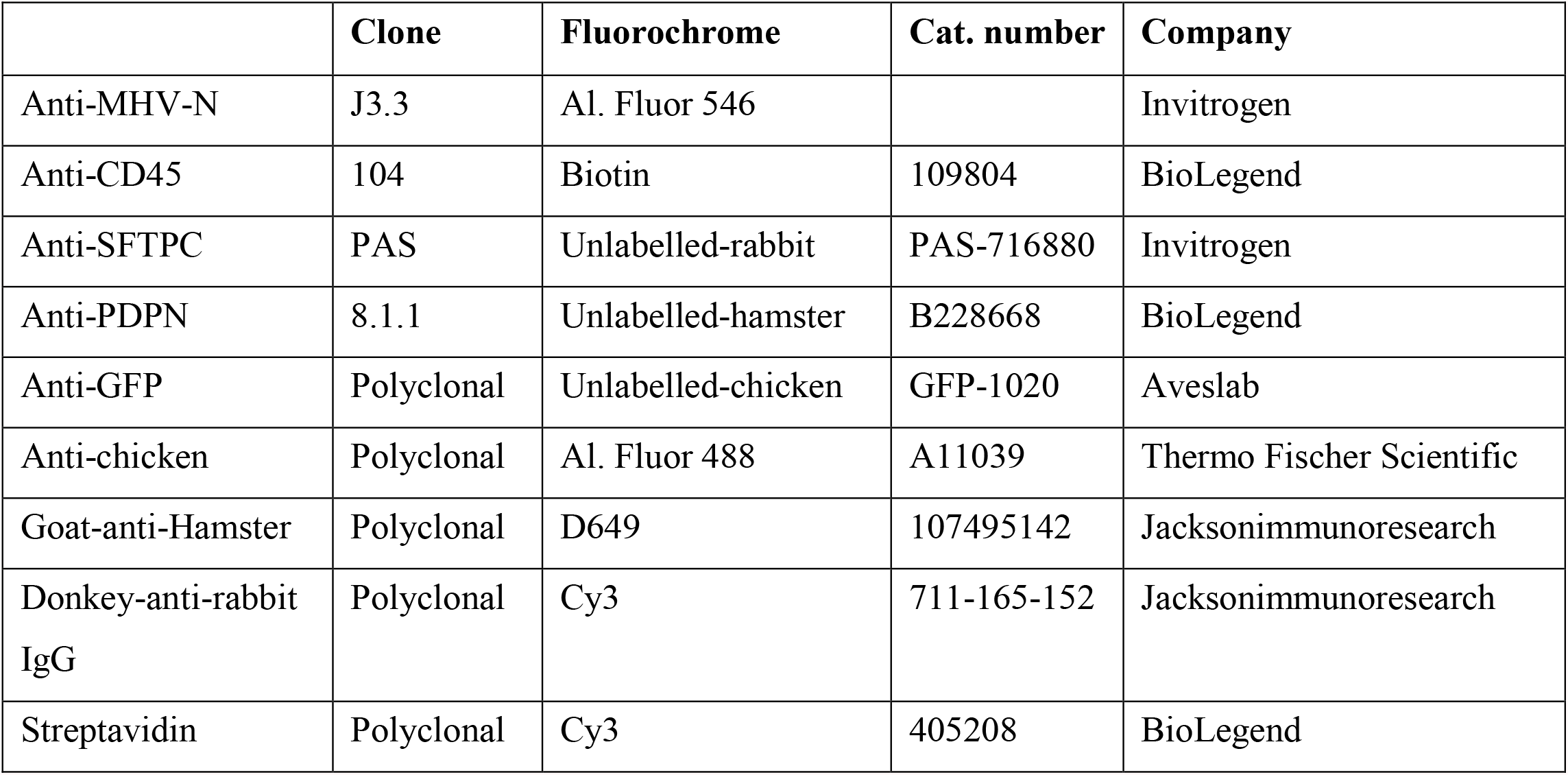
Antibodies for Immunohistochemistry.

|  | **Clone** | **Fluorochrome** | **Cat. number** | **Company** |
| --- | --- | --- | --- | --- |
| Anti-MHV-N | J3.3 | Al. Fluor 546 |  | Invitrogen |
| Anti-CD45 | 104 | Biotin | 109804 | BioLegend |
| Anti-SFTPC | PAS | Unlabelled-rabbit | PAS-716880 | Invitrogen |
| Anti-PDPN | 8.1.1 | Unlabelled-hamster | B228668 | BioLegend |
| Anti-GFP | Polyclonal | Unlabelled-chicken | GFP-1020 | Aveslab |
| Anti-chicken | Polyclonal | Al. Fluor 488 | A11039 | Thermo Fischer Scientific |
| Goat-anti-Hamster | Polyclonal | D649 | 107495142 | Jacksonimmunoresearch |
| Donkey-anti-rabbit<br>IgG | Polyclonal | Cy3 | 711-165-152 | Jacksonimmunoresearch |
| Streptavidin | Polyclonal | Cy3 | 405208 | BioLegend |

