## Supplementary Figures for "TYPE I INTERFERON-DEPENDENT CELLULAR REMODELING DURING ACUTE PULMONARY MHV INFECTION"

#### SUPPLEMENTARY DATA

**SUPPLEMENTARY FIGURE LEGENDS**

**SUPPLEMENTARY FIGURE 1**

A) Infection status of mice in nasal associated lymphoid tissue (NALT), trachea, lung and cervical LN (cLN) over time in mice infected intranasally with $5x10^4$ PFU MHV. Blue color demarcates mice without detectable viral titers in the respective organ. B) Gating strategy of myeloid cell populations in mouse lungs

C) Representative flow cytometric plots of inflammatory macrophages and interstitial macrophages over time in $5x10^4$ PFU MHV infected B6 mice. D) Gating strategy of $s598^+$ $CD8^+$ T cells and E) the respective representative flow cytometric plots of MHV-infected B6 mouse lungs at the indicated time points F) Gating strategy of $KLRG1^+$ $CD62L^-$ $CD8^+$ T cells at the indicated days after infection and G) respective representative flow cytometric plots over time.

**SUPPLEMENTARY FIGURE 2**

A) Marker genes of naïve lung populations B) Gating strategy to identify stromal and hematopoietic cell populations. C-D) Relative composition of MHV-GFP infected cell types within the C) $CD45^-$ $GFP^+$ and D) $CD45^+$ $GFP^+$ populations at 48 hours post infection.

**SUPPLEMENTARY FIGURE 3**

A-C) Representative flow cytometric plots of A) inflammatory macrophages and interstitial macrophages B) alveolar macrophages and C) neutrophils in infected LysM-Cre *Ifnar*$^{fl/fl}$ and LMC at 2 or 4 days post infection.

### Supplementary Figure 1

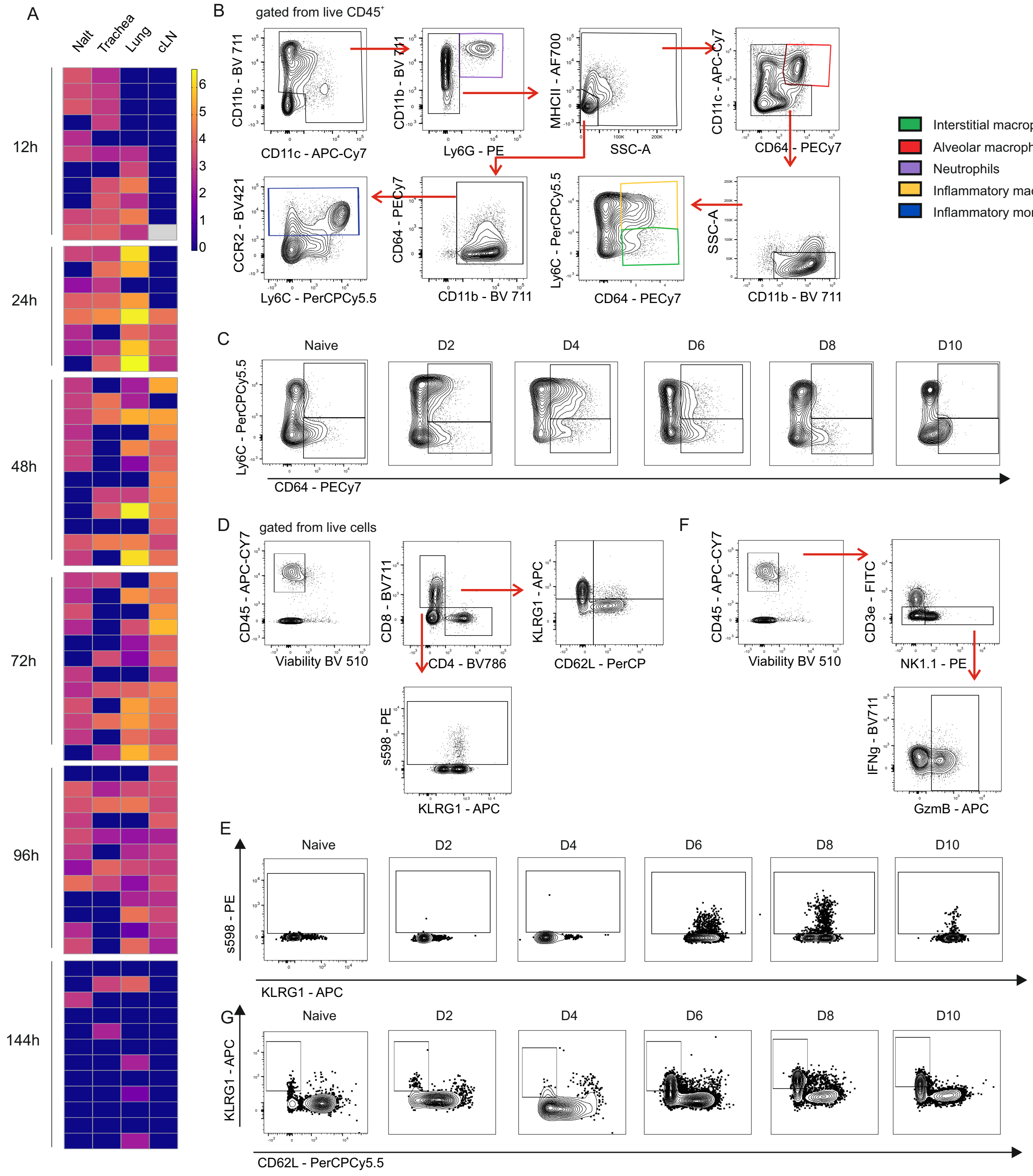

### Supplementary Figure 2

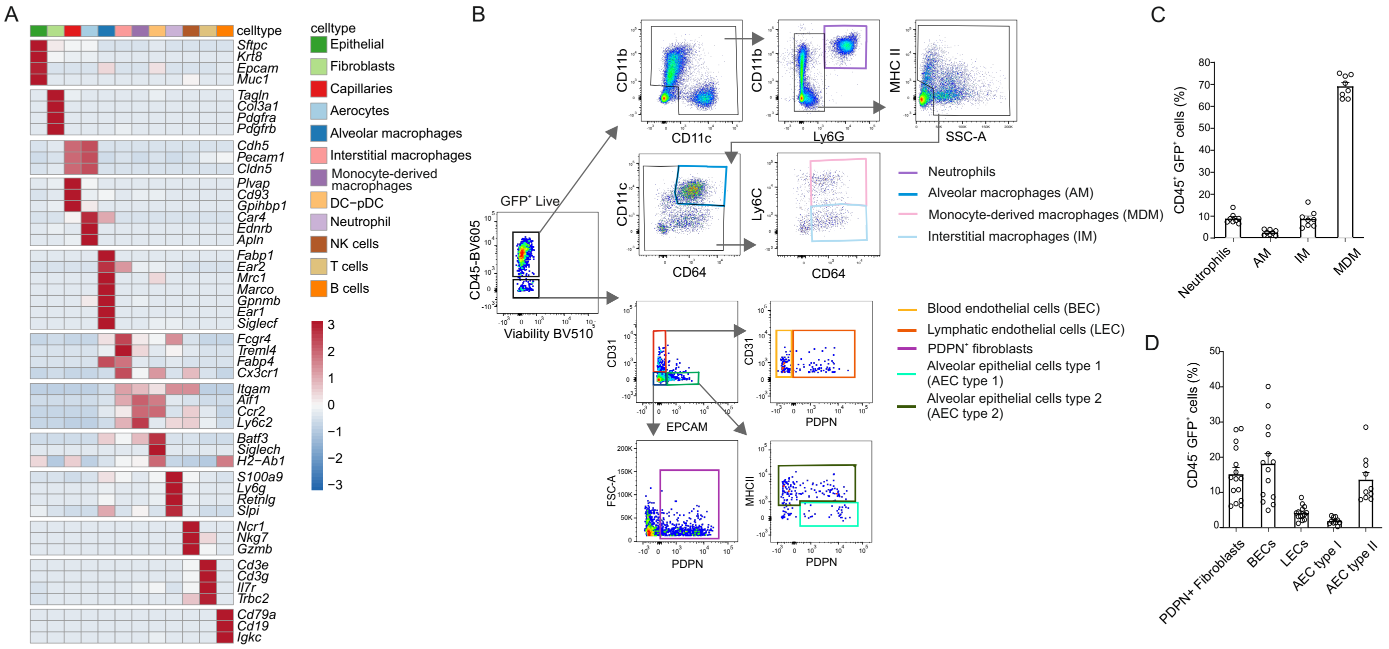

### Supplementary Figure 3

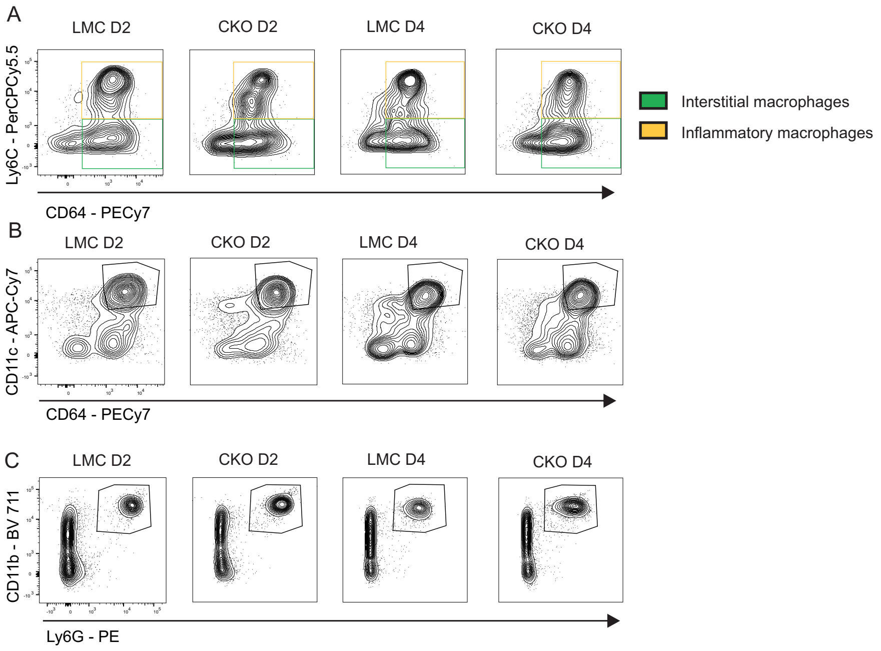
